# Behavioral hierarchy without a hierarchical brain

**DOI:** 10.64898/2026.03.01.708904

**Authors:** Yaning Han, Quanying Liu, Siyuan Liu, Heping Cheng, Pengfei Wei

## Abstract

Behavior is inherently hierarchical. The prevailing view holds that hierarchical behavior arises from the brain’s hierarchical organization. However, this view is primarily derived from experiments in highly controlled laboratory settings, and it remains unknown whether the same principles apply during natural, freely moving behavior. Here, we show that, under naturalistic and freely moving conditions, hierarchical mouse behavior can emerge from localized cortical dynamics alone, independent of anatomical or functional brain hierarchies. We first established a naturalistic neuroethological platform that enables quantitative characterization of hierarchical behavior alongside its corresponding neural activity. Using this platform, we find that localized cortical dynamics encode multiple levels of behavioral hierarchy. Specifically, low-dimensional neural dynamics encode high-level behavioral composites, whereas higher-dimensional neural dynamics encode low-level behavioral kinematics. Finally, by selectively perturbing high-dimensional components of cortical dynamics using optogenetics, we observe a selective reduction in low-level kinematic behaviors, providing causal evidence for this relationship. Together, these findings challenge a central assumption in systems neuroscience by demonstrating that hierarchical behavior does not require a hierarchically organized brain. Instead, hierarchy emerges from the dimensional organization of localized cortical dynamics, revealing a previously unrecognized principle by which complex behavior can arise from relatively simple neural substrates. This dynamic principle offers a unifying framework for understanding how biological systems achieve behavioral flexibility under natural conditions and suggests new directions for artificial intelligence that rely on adaptive local dynamics rather than increasingly deep architectures.

## Introduction

Survival depends on the hierarchical organization of behavior^1^. From humans to other animals, complex behaviors can be decomposed into multiple levels, including high-level behavioral goals, intermediate behavioral units, and low-level motor control^2-4^. For example, during social interactions in mice, high-hierarchy goals involve initiating or maintaining social engagement, intermediate hierarchy consists of reusable behavioral units such as approach, following, sniffing, and avoidance, and low-hierarchy execution relies on continuous postural adjustments and fine motor control^5,6^. These behavioral hierarchies are temporally nested and dynamically interleaved, together giving rise to coherent and flexible natural behavior^7^. However, how the brain supports and implements such a hierarchical organization of behavior remains an ongoing debate.

A widely adopted interpretative framework proposes that hierarchical behavioral organization corresponds to hierarchical organization in the brain^4^. For example, studies in humans and other animals performing sequence learning tasks have reported differential neural representations associated with distinct hierarchies of behavioral structure^8-10^. These investigations of hierarchical neural representations of behavior have relied on simplified experimental paradigms, such as head-fixed preparations or discrete binary sequential decision tasks. While these paradigms allow for precise experimental control, they constrain motor freedom and behavioral diversity^11^. These experimental controls limit the range of neural dynamical states expressed by the brain and, consequently, influence inferences about the organization of behavioral hierarchies^12^.

Studies in freely moving primates have shown that neural activity under naturalistic conditions significantly differs from that observed in head-fixed settings. Even during similar behaviors, neural responses exhibit substantially greater variability^13^. Behavior-related signals can be distributed across cortical regions traditionally associated primarily with visual processing^14^. In parallel, theoretical work such as the “shallow brain” hypothesis has proposed that relatively non-hierarchical local neural microcircuits possess sufficient computational capacity to support complex information processing and behavior, further suggesting that behavioral hierarchies need not map directly onto anatomical hierarchies^15^.

To answer whether the hierarchical organization of natural behavior corresponds to a hierarchical organization of the brain, we established a naturalistic neuroethological platform based on the hierarchical behavior quantification methods previously developed in our laboratory^16-18^ with miniature two-photon calcium imaging (mTPM) in freely social mice^19-21^. We found that neural dynamics in the primary somatosensory cortex (S1), dorsomedial prefrontal cortex (dmPFC), and primary motor cortex (M1) contain temporally distinct dynamical dimensions that simultaneously encode different hierarchies of behavior. These cortical dynamics are structurally embedded within a shared low-dimensional manifold across mice and their brain areas. Behavioral hierarchy is therefore not segregated into specific brain regions. Instead, different behavioral hierarchies are expressed in parallel through distributed dynamical modes across multiple localized cortical areas.

To establish a causal link between localized cortical dynamics and hierarchical behavior, we perturbed the dmPFC using optogenetics. This perturbation increased dynamical noise in the high-dimensional localized cortical dynamics but not their low-dimensional counterparts. Correspondingly, high-hierarchy behavioral motifs remained unchanged, whereas the fine-grained, low-hierarchical pose kinematics were significantly reduced. These results provide causal evidence that distinct dimensions of localized cortical dynamics make separable functional contributions to different behavioral hierarchies.

These findings support a new view in which hierarchical organization of natural behavior does not necessarily follow a strict anatomical or functional hierarchy, but instead emerges from the conserved organization of local dynamics within different cortical networks. This study proposes a generalizable neural computational principle, whereby natural hierarchical behavioral structure can be realized through the localized cortical dynamics across dimensions and timescales, without requiring a pre-specified structural hierarchy of the brain.

## Results

### A naturalistic neuroethological platform

Building on our previous work in resolving multiscale animal behavior, we developed an integrated neuroethological platform that models hierarchical behavior and records neural activity in freely moving mice (Fig. 1B-F). The core of this platform is MouseVenue3D^21,22^, which integrates a miniature two-photon microscope (mTPM)^19,20^ with a synchronized multi-view camera array, enabling simultaneous recording of hundreds of neurons together with high-resolution behavior videos from freely moving mice (Fig. 1A). In this study, we implemented naturalistic social interaction by allowing a subject mouse carrying the mTPM and an object mouse without imaging equipment to interact in an open-field arena freely.

**Fig. 1.**
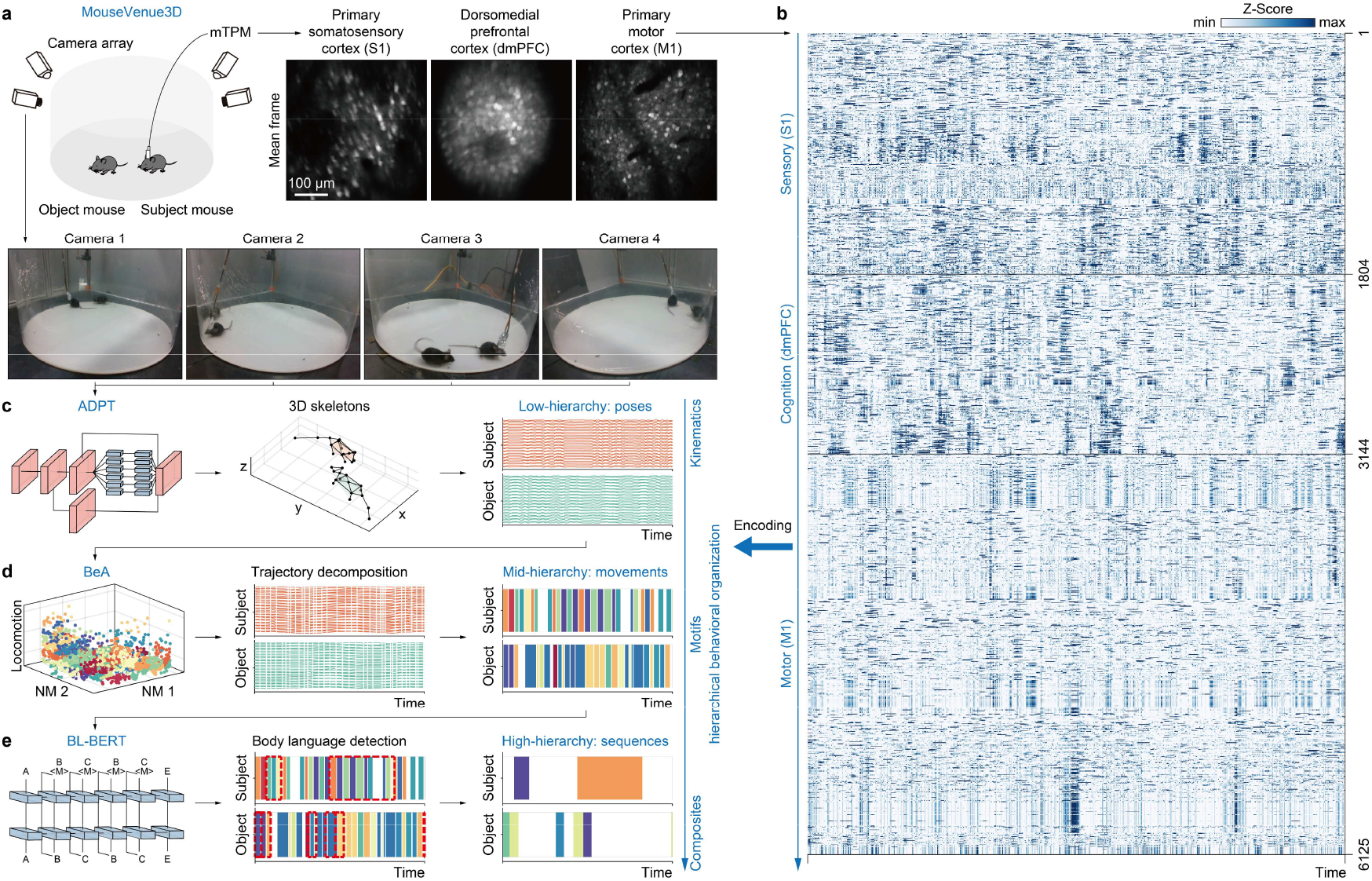
An integrated neuroethological platform modeling hierarchical behavior for naturalistic neural decoding. **a**, MouseVenue3D system for simultaneous multi-view behavior capture and two-photon microscopy (mTPM) neural imaging. A camera array records the subject and object mice from four synchronized views, while brain activity is imaged in separate experiments from the primary somatosensory cortex (S1), the dorsomedial prefrontal cortex (dmPFC), and the primary motor cortex (M1). In all experiments, the mouse implanted for mTPM imaging is referred to as the subject mouse, whereas the unimplanted mouse is referred to as the object mouse. **b**, Neural population activity heatmaps from S1, dmPFC, and M1 (n = 6,125 neurons) from 15 mice. **c**, Low-hierarchy pose kinematics were extracted using the Anti-Drift Pose Tracker (ADPT)^16^, which combines 2D pose estimation and 3D keypoint reconstructions. **d**, Mid-hierarchy movement motifs were decomposed by unsupervised clustering of locomotor dynamics using the Behavior Atlas (BeA)^17^. **e**, High-hierarchy behavioral sequence composites were derived using BL-BERT^18^, a transformer-based sequence model that identifies body-language structures. Red boxes mark automatically detected segment boundaries. The overlaid hierarchical behavioral labels (right, from **c** to **e**) illustrate the progression from low-level kinematics to mid-level motifs and high-level composites, providing a structured basis for investigating cortical encoding of behavior.

Because social interaction requires sensory perception of a partner, cognitive evaluation of social cues, and motor control of one’s own actions, we targeted three key brain regions accordingly: the S1 for sensory monitoring, the dmPFC for cognitive monitoring, and the M1 for motor monitoring. These regions were recorded in 15 mice in total, with 5 mice per brain area. After neural signal extraction, we obtained 1,804 neurons in S1, 1,340 neurons in dmPFC, and 2,981 neurons in M1 for subsequent brain-behavior mapping analyses (Fig. 1B).

We constructed a hierarchical representation of behavior by sequentially applying three computational models. First, the Anti-Drift Pose Tracker (ADPT)^16,23^ provided stable, frame-by-frame 3D keypoints that defined the pose hierarchy (Fig. 1D). Next, we used the Behavior Atlas (BeA) framework^17,24^ to segment continuous posture into discrete movement fragments based on posture-defined transitions (Fig. 1E). Finally, we applied the Body Language Bidirectional Encoder Representation from Transformers (BL-BERT)^18,25^, a self-supervised deep learning model to identify behavioral sequences from these movements (Fig. 1F). These pose, movement, and sequence, form a hierarchical organization of behavior with increasing temporal scale from kinematics to composites, allowing us to examine how different cortical areas represent behavior across these levels.

This naturalistic neuroethological platform provides a unified framework for linking hierarchical behavioral organization with large-scale cortical activity during naturalistic social interaction. By combining a hierarchical behavioral quantification framework and brain mTPM imaging across sensory, cognitive, and motor circuits, this platform establishes the conditions to investigate how neural activity encodes behavioral features at different hierarchies.

### Localized cortical dynamics reflect hierarchical behavior

To obtain an ethologically grounded baseline, we first focused on social interaction episodes whose stereotyped far-close-far phases provided an internal within-trial reference for cross-trial alignment. Social interactions comprise a close phase involving body interaction, and a far phase reflecting independent behavior (Fig. 2A). The corresponding subject and object movements aligned to the interaction onset and offset (Fig. 2B). Nevertheless, the single-neuron activity appeared disordered (Fig. 2C). Despite apparent disorder at the single-neuron level, the normalized PCs show aligned dynamic patterns with the social interaction episode (Fig. 2D). As the index of the PCs increases, the aligned patterns become less clear.

**Fig. 2.**
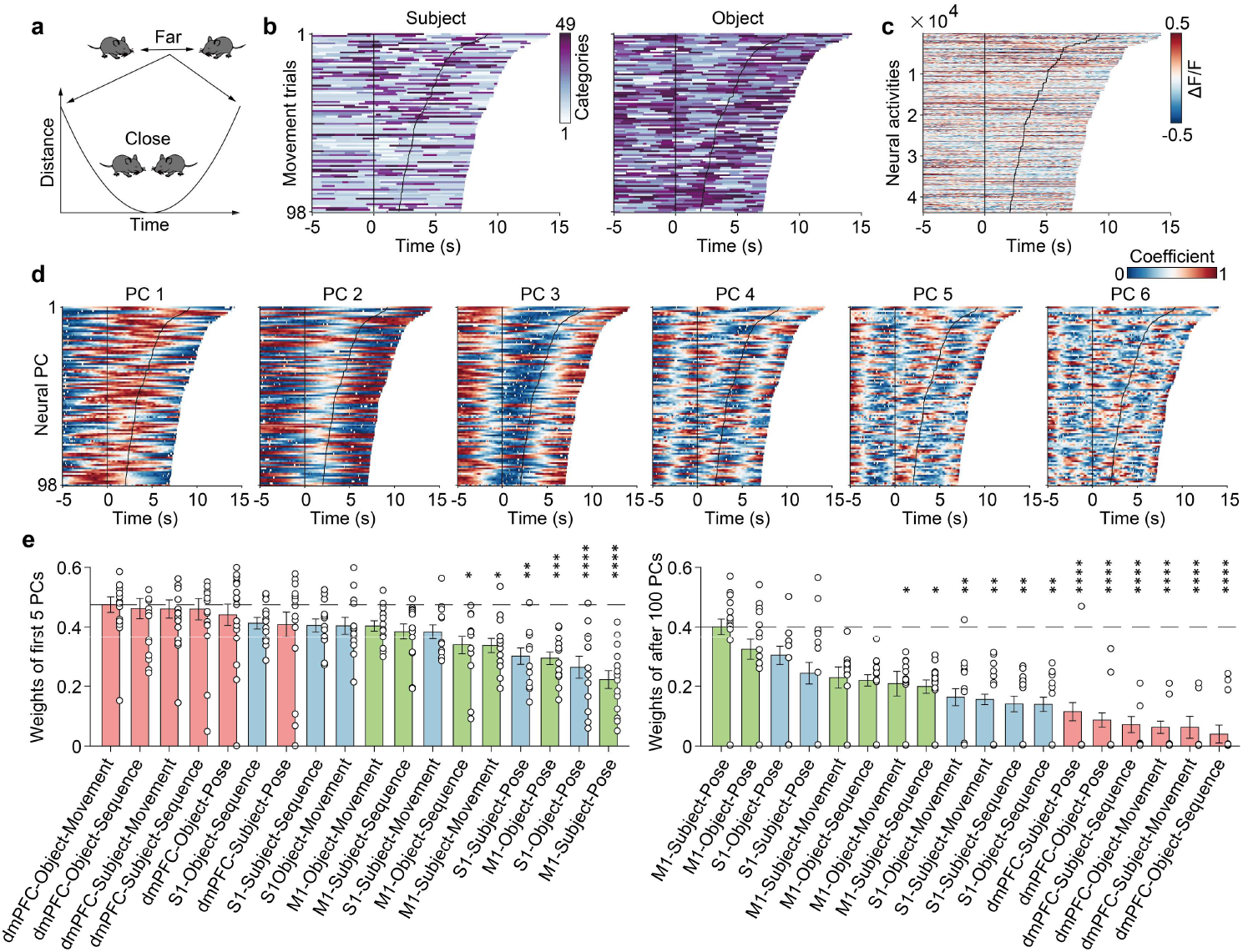
Localized cortical dynamics encode hierarchical behavior through multiple dimensions. **a**, Social interaction episodes segmented based on distance between two mice (far → close → far). **b**, Discrete movement series of subject and object mice during social interaction episodes, sorted by trial duration. Each trial includes 5 s of baseline (far), dynamic close interaction (close), and 5 s of post-interaction baseline (far). **c**, Activity of each neuron during the episodes shown in **b**. **d**, First six principal components (PCs) extracted from trial-wise neural activity in **c**, revealing alignment with social interaction episodes. **e**, Linear decoding weights predicting hierarchical behavioral organizations from neural PCs. Left: weights from first five PCs (One-way ANOVA followed by Dunnett’s multiple comparisons test, mean±SEM, N=18, 18, 18, 18, 18, 13, 18, 13, 13,14,14, 13,14,14, 13,14,13, and 14, the P values from left to right are >0.9999, >0.9999, >0.9999, 0.9972, 0.8142, 0.6305, 0.6814, 0.6618, 0.6248, 0.3000, 0.3105, 0.0190, 0.0152, 0.0010, 0.0004, <0.0001, and <0.0001). Right: weights from PCs beyond the 100th (One-way ANOVA followed by Dunnett’s multiple comparisons test, mean±SEM, N=14, 14, 13, 13, 14, 14, 14, 14, 13, 13, 13, 13, 18, 18, 18, 18, 18, and 18, the P values from left to right are 0.9158, 0.7339, 0.1593, 0.0847, 0.0583, 0.0365, 0.0236, 0.0049, 0.0033, 0.0014, 0.0013, <0.0001, <0.0001, <0.0001, <0.0001, <0.0001, and <0.0001). *P < 0.05, **P < 0.01, ***P < 0.001, ****P < 0.0001.

**Fig. 3.**
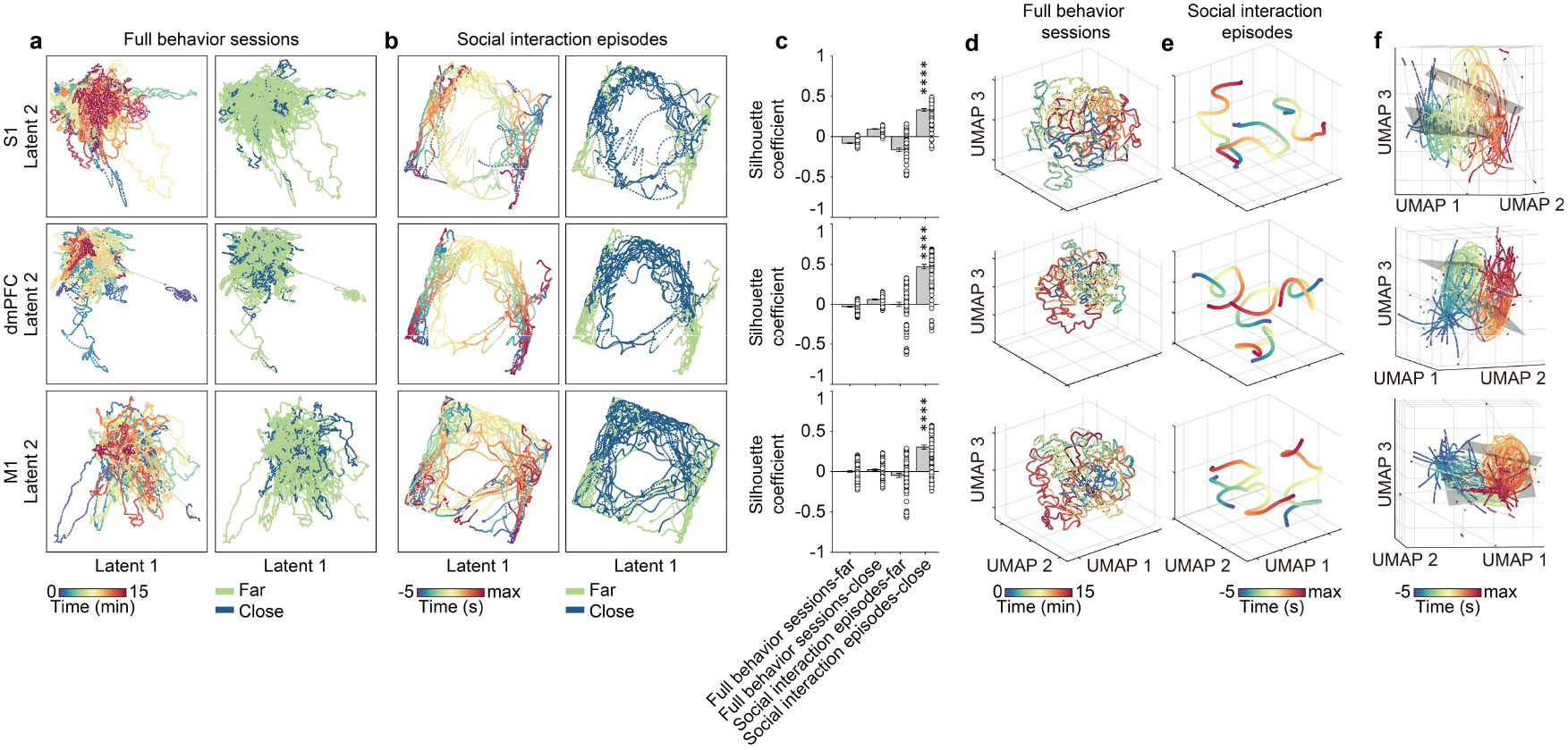
Localized cortical dynamics are conserved across trials, brain areas, and mice. **a**, rSLDS-derived latent embeddings of neural PCs in S1, dmPFC, and M1 during full behavior sessions. Left: embeddings colored by time. Right: embeddings colored by distance (far vs. close). **b**, rSLDS-derived latent embeddings during social interaction episodes. **c**, Silhouette coefficients quantifying cluster separability in **a** and **b** (One-way ANOVA followed by Tukey multiple comparison test, mean±SEM, N=100, P values between social interaction episodes and other groups are all <0.0001). **d**, UMAP embeddings of neural activity from S1, dmPFC, and M1 during full behavior sessions. **e**, UMAP embeddings of neural activity during social interaction episodes. **f**, GTW-aligned UMAP trajectories of neural dynamics during social interaction episodes across S1, dmPFC, and M1. ****P < 0.0001.

To relate neural PCs to hierarchical behavioral organization, we fit linear models predicting pose, movement, and sequence from neural PCs in S1, M1, and dmPFC across subject and object mice (Fig. 2E). PCs 1 to 5 weighted “dmPFC”, “movement”, and “sequence”, whereas PCs > 100 weighted “S1”, “M1”, and “pose”, indicating composite content in low order PCs and kinematic features in high order PCs. This conserved mechanism broadly exists in these cortical areas. Thus, local circuits can represent behavioral hierarchies through different weightings of local dynamic dimensions.

These results show that disordered single-neuron activity resolves into structured, multi-dimensional localized cortical dynamics that track social interaction episodes in a temporally aligned manner. Distinct neural PCs differentially relate to multiple levels of behavioral organization: low-order PCs preferentially encode composite, higher-level features such as social engagement and behavioral sequences, whereas higher-order PCs emphasize kinematic details like poses. This graded mapping is conserved across S1, M1, and dmPFC, indicating that hierarchical behavior is not encoded in hierarchical brain areas but is instead represented through differential weightings of dynamic dimensions within localized cortical circuits.

### Localized cortical dynamics share conserved structure

Although the above results revealed that different neural PCs encode behavioral hierarchies, it remains unclear whether these PCs follow consistent population dynamics across trials and across individuals. If such consistency exists, it should manifest as reproducible low-dimensional trajectories that can be captured by a shared dynamical model. To test this possibility, we concatenated the trial-aligned neural PCs and applied a recurrent switching linear dynamical system (rSLDS) ^26-29^, which identifies discrete latent states together with their associated continuous population flow fields (Fig. 2A-B).

Across full behavioral sessions, the rSLDS trajectories exhibited large variability (Fig. 2A), consistent with the diverse and continuously shifting behaviors expressed in naturalistic contexts. In contrast, when restricting the analysis to social interaction episodes, the rSLDS uncovered separable dynamical patterns across trials and across animals between far and close phases (Fig. 2B). To quantify this structure, we computed silhouette coefficients for the latent trajectories (Fig. 2C). The close-phase social interactions showed significantly higher silhouette coefficients than any other condition, illustrating well-separated latent-state clusters. These findings show that neural PC dynamics reorganize flexibly across the full spectrum of natural behaviors but become locally consistent when aligned to ethologically meaningful social cues.

To further compare global neural dynamics with their localized counterparts, we next embedded the neural activities in a common low-dimensional space using UMAP (Fig. 2D, E). When embedding the full behavioral sessions, trajectories from S1, dmPFC, and M1 wandered continuously through state space with no obvious recurring pattern (Fig. 2D). In contrast, when we isolated social interaction episodes and embedded only these segments, we observed trajectories with similar far-close-far shapes across animals and regions, but their absolute positions in the UMAP space differed (Fig. 2E). Thus, cortical dynamics during social episodes appeared to follow a stereotyped pattern, yet were embedded at different locations along the global population flow.

To assess whether these far-close-far structures shared a common underlying geometric organization, we applied Generalized Time Warping (GTW)^30^ to align the trajectories across animals and trials (Fig. 2K). After GTW alignment, the UMAP trajectories converged onto a consistent low-dimensional manifold flow. This revealed a conserved temporal ordering and curvature of localized cortical dynamics, despite differences in absolute embedding positions. These results demonstrate that local cortical dynamics across S1, dmPFC, and M1 share a common transient structure during social interaction episodes, realized as a stereotyped dynamical motif expressed within region-specific manifolds.

Together, these results show that the cortical dynamics follow reproducible latent trajectories and manifolds that can be captured by a shared dynamical model across trials, animals, and cortical areas. This reveals that cortical dynamics are globally continuous and fluid over the full behavioral repertoire, yet transient, locally stabilized, and highly consistent when aligned to ethologically meaningful social events.

### Localized cortical dynamics predict behavior

Building on the analyses focused on social interaction episodes, we next extended our approach to a more general behavioral context. To retain an ethologically grounded reference, we used social interaction episodes as an internal baseline and examined the neuroethology of the intervals between successive interactions, which we term post-interaction behavior (Fig. 4A). This analysis allowed us to test whether localized cortical dynamics generalize beyond the interaction itself.

**Fig. 4.**
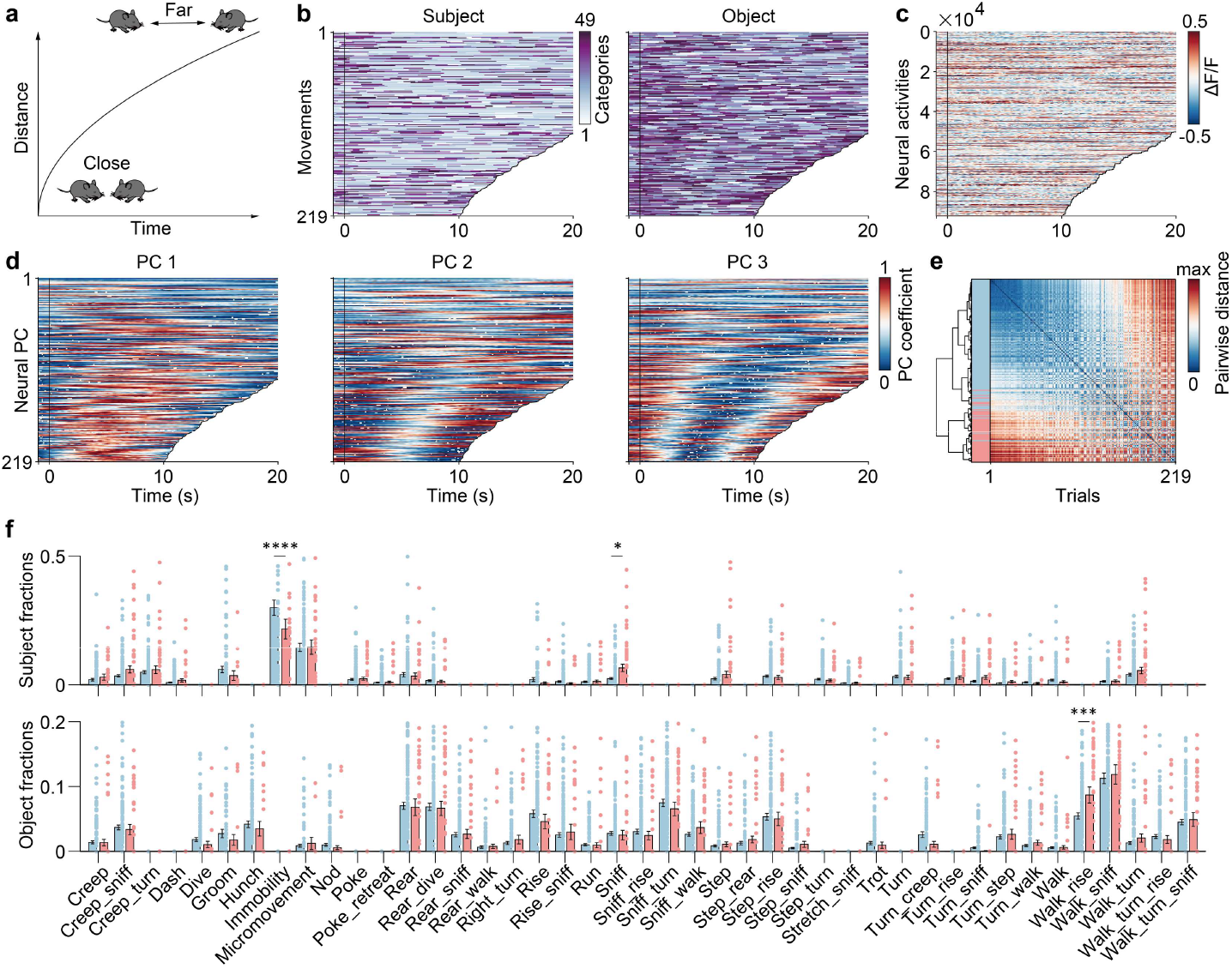
Predictable post-interaction behavior through localized cortical dynamics. **a**, The trials of post-interaction are split for later analysis. The stop point is the start time of the next social interaction. **b**, The movements of the subject and object mice. The trials are sorted by their durations. Each trial includes 1s of close social interaction as a baseline, and the end is before the next close interaction. **c**, Neural activities corresponding to **b**. **d**, The first 3 PCs of neural activities of **c**. **e**, The clustering of pairwise distances of the first 3 PCs. **f**, Movement differences between subject and object mice were compared according to the two clusters of neural PCs (Two-way ANOVA followed by Sidak’s multiple comparisons test, adjusted P values of subject fractions (mean±SEM, N=153) are >0.9999, 0.7408, >0.9999, >0.9999, >0.9999, 0.8489, >0.9999, <0.0001, >0.9999, >0.9999, >0.9999, >0.9999, >0.9999, >0.9999, >0.9999, >0.9999, >0.9999, >0.99 99, >0.9999, >0.9999, 0.0173, >0.9999, >0.9999, >0.9999, 0.9990, >0.9999, >0.9999, >0.9999, >0.9999, >0.9999, >0.9999, >0.9999, >0.9999, >0.9999, >0.999 9, >0.9999, >0.9999, >0.9999, >0.9999, >0.9999, 0.9998, 0.9999, >0.9999. Adjusted P values of object fractions (mean±SEM, N=66) are >0.9999, >0.9999, >0.9999, >0.9999, >0.9999, 0.9995, >0.9999, >0.9999, >0.9999, >0.9999, >0.9999, >0.9999, >0.9999, >0.9999, >0.9999, >0.999 9, >0.9999, 0.9796, >0.9999, >0.9999, >0.9999, >0.9999, 0.9999, 0.9990, >0.9999, >0.9999, >0.9999, >0.9999, >0.9999, >0.9999, >0.9999, >0.9999, 0.7783, >0.9999, >0.9999, >0.9999, >0.9999, >0.9999, 0.0002, >0.9999, >0.9999, >0.9999, >0.9999). *P < 0.05, ***P < 0.001, ****P < 0.0001.

**Fig. 5.**
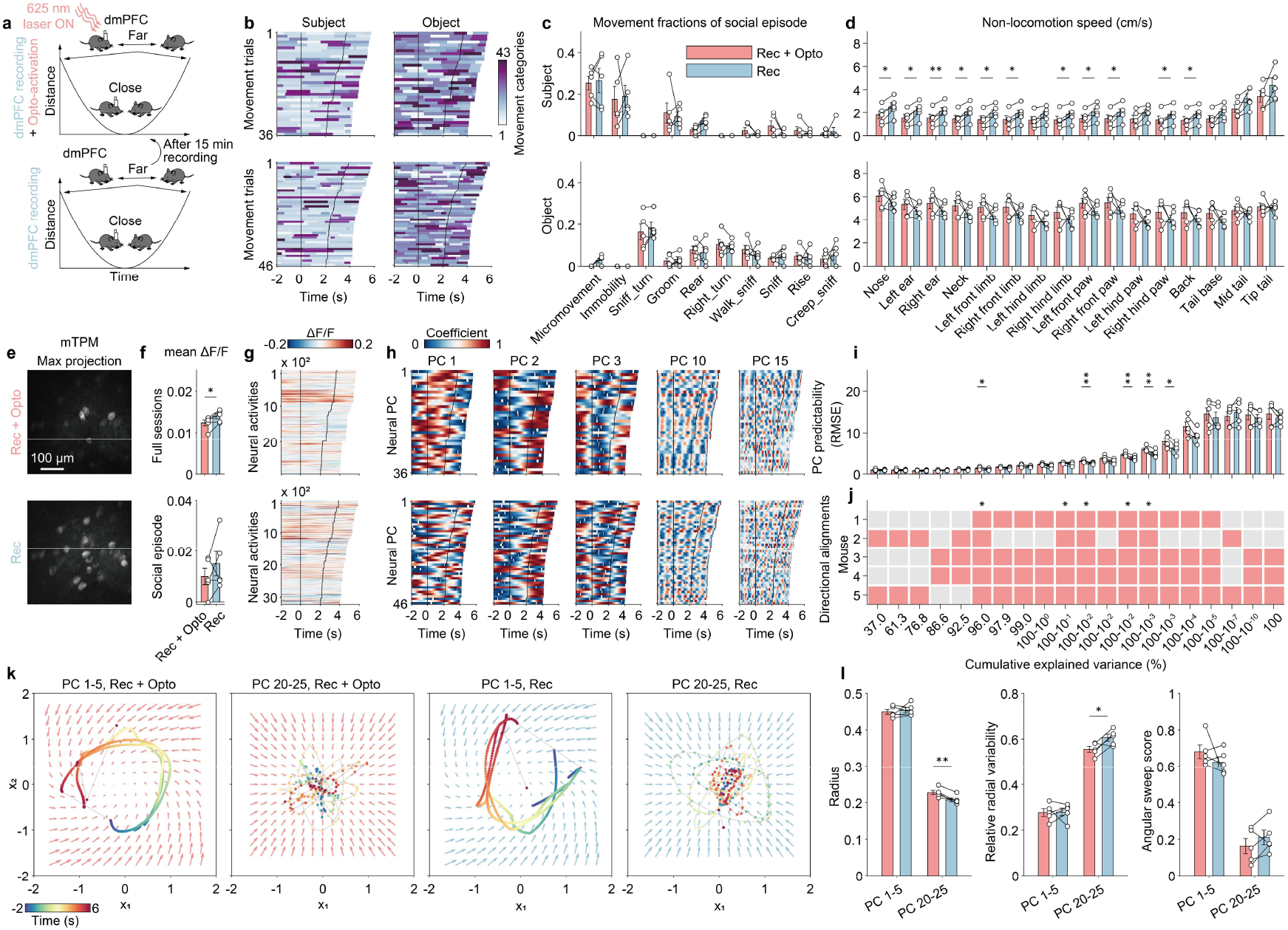
Optogenetic perturbation of dmPFC disrupts high-dimensional neural components and selectively alters hierarchical behaviors. **a**, Social interaction episodes were extracted for comparison between dmPFC optogenetic activation (Rec + Opto, upper) and no-stimulation control (Rec, lower). Each mouse underwent 15 min of recording without stimulation, followed by 15 min of continuous 625-nm activation. For both sessions, social interaction episodes were segmented post hoc from the full recordings based on distance. **b**, Movement of subject and object mice during the extracted social interaction episodes. **c**, Movement fractions during the extracted social interaction episodes for the Rec + Opto and the Rec conditions. All movement fractions showed no significant differences between conditions. Only the top ten categories are shown for visualization (two-way ANOVA followed by Sidak multiple comparisons test, mean±SEM, N=5, the P values of subject movement fractions from left to right are >0.9999, >0.9999, >0.9999, >0.9999, 0.9692, >0.9999, >0.9999, 0.9893, >0.9999, and >0.9999. The P values of object movement fractions from left to right are 0.9926, >0.9999, >0.9999, >0.9999, >0.9999, >0.9999, 0.9918, >0.9999, >0.9999, and >0.9999). **d**, Non-locomotion speed across body parts during the extracted social interaction episodes for the Rec + Opto and Rec conditions. Optogenetic perturbation of dmPFC modulated non-locomotor speeds in specific body parts (paired t-test and sign-rank test, mean±SEM, N=5, the P values of subject non-locomotion speed from left to right are 0.0137, 0.0112, 0.0099, 0.0114, 0.0446, 0.0117, 0.0925, 0.0252, 0.0318, 0.0164, 0.0601, 0.0138, 0.0224, 0.0518, 0.0643, and 0.1334. The P values of object non-locomotion speed from left to right are 0.3914, 0.3857, 0.3797, 0.3715, 0.3005, 0.4303, 0.3987, 0.4523, 0.3100, 0.3568, 0.4278, 0.4401, 0.4813, 0.4009, 0.6369, and 0.8359). **e**, Max-projection mTPM images from dmPFC for the Rec + Opto and Rec conditions, taken from the full behavior sessions. **f**, Mean ΔF/F of dmPFC activity for the Rec + Opto and the Rec conditions. Top: mean ΔF/F across the full 15-min recording sessions (Paired t-test, mean±SEM, N=5, the P value is 0.0184). Bottom: mean ΔF/F computed only within the extracted social interaction episodes (Paired t-test, mean±SEM, N=5, the P value is 0.3015). **g**, Activity of each neuron during the social interaction episodes. **h**, Neural PC 1, 2, 3, 10, and 15 extracted from **g**. **i**, RMSE of PC predictability for the optogenetic activation (Rec + Opto) and control (Rec) conditions (paired t-test, mean±SEM, N=5, the P values from left to right are 0.8679, 0.9616, 0.4299, 0.9701, 0.3125, 0.0409, 0.1296, 0.1525, 0.1189, 0.0625, 0.0048, 0.0857, 0.0091, 0.0064, 0.0560, 0.4099, 0.2210, 0.3415, and 0.2567). **j**, Directional alignments between non-locomotion speed and PC predictability in each mouse (one-sided exact binomial test, N=5, P values from left to right are 0.8125, 0.8125, 0.8125, 0.8125, 0.5000, 0.0312, 0.1875, 0.1875, 0.1875, 0.0312, 0.0312, 0.1875, 0.0312, 0.0312, 0.1875, 0.1875, 0.5000, 0.8125, 0.5000, and 0.5000). **k**, Flow fields reconstructed from low-dimensional (PCs 1-5) and high-dimensional (PCs 20-25) neural PCs, modeled using rSLDS for the Rec + Opto and Rec conditions. **l**, Quantification of flow field structure in **k**. Left: Radius, reflecting the spatial extent of neural state trajectories (paired t-test, mean±SEM, N=5, P values are 0.6450 and 0.0064). Middle: Relative radial variability, quantifying the heterogeneity of radial components in the flow field (paired t-test, mean±SEM, N=5, P values are 0.9270 and 0.0379). Right: Angular sweep score, measuring the directional coverage and rotational organization of neural dynamics (paired t-test, mean±SEM, N=5, P values are 0.3439 and 0.2257). *P < 0.05, **P < 0.01, ***P < 0.001, ****P < 0.0001.

Post-interaction behavior exhibited substantially more diverse and complex action patterns compared to the interaction phase (Fig. 4B). As during social interaction, single-neuron activity did not reveal stereotyped structure (Fig. 4C). In contrast, neural PCs displayed clear and reproducible dynamic patterns (Fig. 4D). Notably, these patterns differed between short-duration (<20 s) and long-duration (>20 s) post-interaction periods, suggesting the presence of distinct underlying behavioral states.

To determine whether these distinct neural patterns corresponded to different behavioral outcomes, we applied an unsupervised analysis using a Dynamic Time Alignment Kernel (DTAK)^17,24^ combined with hierarchical clustering (Fig. 4E). DTAK aligns intrinsic structural similarities within time series while remaining insensitive to differences in absolute duration. For descriptive clarity, we refer to the resulting clusters as a short-duration group (red) and a long-duration group (blue).

Based on the unsupervised clustering, we partitioned the corresponding behavioral segments and quantified movement fractions with manual annotations, separately for the subject and object mice (Fig. 4F). In the long-duration group, subject mice exhibited significantly higher immobility compared to the short-duration group. In contrast, subject mice in the short-duration group showed increased sniffing, accompanied by elevated walk-rise behavior in the object mice. These patterns predict two distinct post-interaction behavioral phenotypes. The short-duration cluster corresponds to episodes in which mice re-initiate social interaction within approximately 10 seconds, characterized by preparatory behaviors such as sniffing and walk-rise. In contrast, the long-duration cluster reflects disengagement from social interaction, with the subject mouse remaining largely immobile and not preparing for re-engagement.

These results demonstrate that localized cortical dynamics not only reflect ongoing social interaction but also predict subsequent behavioral phenotypes beyond the interaction itself. Despite the absence of interpretable structure at the single-neuron level, low-dimensional cortical dynamics reliably separate post-interaction periods into distinct states that forecast whether animals re-engage in social behavior or disengage. This predictive organization generalizes across variable temporal scales and hierarchical behavioral contexts.

### Perturbing localized cortical dynamics reshapes hierarchical behavior

Finally, to test the causality from localized cortical dynamics to hierarchical behavior, we used optogenetics to perturb the dmPFC of mice (Fig. 3A). Because the dmPFC is a key hub for social behavior ^5,31^, we selected it as the target region for perturbation. We applied the same naturalistic social-interaction paradigm as in the previous experiments. Neural activity and behavior were first recorded for 15 min without stimulation (Fig. 3A, bottom), after which the optogenetic light was switched on to globally activate dmPFC neurons for an additional 15 min (Fig. 3A, top). Social interaction episodes from these recordings were then segmented for further analysis.

The corresponding hierarchical behaviors of the subject and object mice were extracted as before (Fig. 3B). Data shows no significant differences in movement categories between the recording + optogenetic (Rec + Opto) and recording-only (Rec) groups for either the subject or object mice during social episodes (Fig. 3C). This lack of differences at the movement level demonstrates that overall social engagement and the movement hierarchy remain preserved under dmPFC perturbation. We therefore asked whether this optogenetic stimulation might differentially affect specific levels of behavioral organization. A significant change emerged only in the non-locomotion speed of the subject mice, but not the object mice (Fig. 3D). Non-locomotion speed quantifies fine-scale pose kinematics while excluding confounds from high-velocity locomotion. These results demonstrate that this dmPFC perturbation selectively disrupts detailed pose kinematics rather than higher-hierarchy behavioral motifs.

After characterizing the behavioral effects, we next examined how dmPFC neural activity was altered by this perturbation. To assess whether and how neurons were modulated, we visualized the maximum projections of the mTPM recordings (Fig. 3E). Although the opsin is excitatory, global optogenetic activation produced a net suppressive effect at the population level. Approximately 10% of neurons exhibited sustained activation, whereas the remaining ∼90% showed prolonged suppression. Accordingly, the mean ΔF/F across the entire recording session was significantly reduced during optogenetic stimulation compared to the non-stimulation period (Fig. 3F, top). However, when restricting the comparison to social interaction episodes, the mean ΔF/F did not differ between the two conditions (Fig. 3F, bottom). Despite a global perturbation, the dmPFC network reorganizes its population dynamics over time, allowing transient restoration of overall activity levels during social behavior episodes.

We next focused specifically on the social interaction episodes. At the single-neuron level, activity remained heterogeneous in either the Rec + Opto or Rec groups (Fig. 3G). In contrast, as in earlier analyses, neural PCs revealed robust and stereotyped population-level structure across trials and across individuals in both conditions (Fig. 3H). Higher-order PCs displayed increasingly complex temporal structure.

To assess whether this optogenetic perturbation disrupts the multi-dimensional dynamical stability, we quantified the trial-wise predictability of each neural PC using a minimal first-order autoregressive model (Fig. 3I). This optogenetic perturbation significantly increased prediction error for several high-dimensional PCs beyond the 96% cumulative explained variance. It demonstrates that dmPFC activation introduces dynamical noise into high-dimensional components, selectively disturbing their stationarity.

We next asked whether these selective destabilizations were behaviorally relevant. We further examined the directional consistency between optogenetic-induced changes in neural PC predictability and changes in non-locomotion speed on a per-mouse basis (Fig. 3J). Directional alignment was significantly enriched in the same high-dimensional regime in which predictability was degraded. These results provide neuroethological evidence that this optogenetic perturbation induces coordinated reconfiguration of high-dimensional cortical dynamics and fine-grained behavior, with consistent directional coupling between them.

Lastly, we examined whether the localized cortical dynamics of the neural PCs were altered by the perturbation. Because the effects of optogenetics were dimension-dependent, we separately fit rSLDS models to low-dimensional PCs (PCs 1-5) and high-dimensional PCs (PCs 20-25) (Fig. 3K). The resulting flow fields revealed that low-dimensional dynamics were nearly identical between the Rec + Opto and Rec groups, whereas high-dimensional dynamics in the Rec + Opto group exhibited a visibly altered dynamical regime. Quantitatively, the radius of high-dimensional dynamics was significantly larger in the Rec + Opto group than in the Rec group (Fig. 3L, left), while relative radial variability was significantly reduced (Fig. 3L, middle). In contrast, the angular sweep score showed no significant difference between groups (Fig. 3L, right). These results indicate that optogenetic perturbation rescaled the amplitude and stability of localized cortical dynamics without altering their intrinsic phase structure. This selective effect on high-dimensional components corresponds to changes in fine-grained non-locomotion speed and reduced PC predictability. Together, these findings demonstrate that this optogenetic perturbation primarily modulates high-dimensional neuroethological details while preserving the low-dimensional dynamical scaffold underlying natural social behavior.

These results establish a causal pathway from localized cortical dynamics to hierarchical behavior. Although global dmPFC optogenetic perturbation produces relatively homogeneous inhibitory effects at the single-neuron level, it reorganizes neural dynamics at the population level by selectively injecting noise into high-dimensional neural PCs. The resulting changes in high-dimensional neural PCs closely align with perturbations in fine-grained behavioral kinematics, whereas low-dimensional neural PCs remain largely preserved, consistent with the stability of higher-hierarchy movements. This differential perturbation effect provides evidence that multi-dimensional localized cortical dynamics are causally coupled to hierarchical behavioral organization. In particular, perturbations at the neuronal level propagate selectively through high-dimensional neural PCs to shape fine-scale behavior while preserving behavioral compositions.

## Discussion

Whether hierarchical behavior is encoded by hierarchical brain organization remains a longstanding and unresolved question in neuroscience^7^. Using a naturalistic neuroethological platform developed in this study, we systematically examined how sensory, decision-related, and motor cortical areas encode hierarchical behavior in freely interacting mice. Our analyses spanned single-neuron activity, population-level representations, and low-dimensional cortical dynamics, and were complemented by optogenetic perturbations to establish causal links. Across S1, dmPFC, and M1, we found that local cortical circuits generate similar multidimensional dynamics that encode different levels of behavioral hierarchy. Low-dimensional components preferentially represent composite and higher-order behavioral structure, whereas higher-dimensional components capture fine-grained kinematic features. Importantly, optogenetic perturbations at the neuronal level propagated through localized cortical dynamics and selectively altered distinct behavioral levels. These findings provide evidence that hierarchical behavior emerges from localized cortical dynamics spread across regions, rather than being imposed by a strict anatomical or functional hierarchy of the brain. The localized cortical dynamics support a central premise of the “shallow brain” hypothesis that local cortical microcircuits possess sufficient representational capacity to encode complex hierarchical information^15^. This encoding does not rely on an explicit hierarchical mapping across cortical areas. Instead, it emerges from the organization of local neural populations within multi-dimensional dynamical spaces.

Neural dynamics have received increasing attention in recent years, including studies of rotational dynamics^32,33^, attractor states^26-28^, and metastability^34-36^. From a broader dynamical perspective, our findings are conceptually aligned with recent theories of metastability, which emphasize that neural systems dynamically transition among multiple transient quasi-stable states rather than converging to a single stable attractor^34^. Our study further demonstrates that such metastable organization can be instantiated at the level of local cortical microcircuits. Specifically, we show that localized cortical dynamics embedded within a shared low-dimensional neural manifold can directly support the encoding of complex, hierarchical behavior. To distinguish this local form of dynamical organization, we refer to it as localized mesostasis. This term emphasizes a non-equilibrium, metastable dynamical state maintained by local microcircuits within an otherwise established system architecture, enabling flexible and compositional behavioral expression.

Localized mesostasis also offers insights for the design of next-generation artificial intelligence systems. Historically, artificial neural networks have been strongly inspired by the hierarchical organization of the brain, with advances from deep learning to large language models relying on increasingly deep network architectures to enhance representational capacity and task performance^37-39^. While such depth has proven effective for perception and inference, it remains challenging for these architectures to support robust, real-time interaction with dynamic environments, as required in closed-loop control tasks such as adaptive robotics^40^. In contrast, the localized mesostasis observed in natural hierarchical behavior highlights the computational role of multi-dimensional, metastable dynamics within local microcircuits, whose temporal organization and efficiency are well matched to the demands of real-world interaction. Our findings suggest that enhancing the intrinsic dynamical capabilities of individual layers or local modules may provide a complementary route toward improving adaptability and intelligence in artificial systems operating in natural environments.

## Methods

### Animals

Forty male C57BL/6J mice (10-14 weeks old) were used in the experiment. Twenty mice without surgery were housed at 5 mice per cage, and the other fifteen mice with surgery were housed at 1 mouse per cage under a 12 h light-dark cycle at 22–25 °C with 40–70% humidity, and were allowed to access water and food ad libitum (15 in Shenzhen Institutes of Advanced Technology, Shenzhen, China, and another 5 in Nanjing Raygen Health, Nanjing, China). Fifteen mice that underwent surgery were anesthetized with isoflurane and placed in a stereotactic apparatus (RWD). The skull was exposed, and a craniotomy was performed for virus injection. To cover the entire primary somatosensory cortex (S1), dorsomedial prefrontal cortex (dmPFC), and primary motor cortex (M1), the mouse was microinjected at the following coordinates: AP, −0.60mm; ML, −2.40mm; DV, 2.00mm, AP, 2.00mm; ML, −0.30mm; DV, −1.80mm, and AP, 0.30mm; ML, −1.50mm; DV, −0.50mm. AAV9-CaMKII-GCaMP6s viruses were used for S1, and AAV2/9-hsyn-GCaMP6f viruses were used for dmPFC and M1. Five mice used for the optogenetic experiment followed the same surgery procedure as the above dmPFC mice, only the virus was different, which was AAV2/9-hsyn-GCaMP6f and AAV2/9-hsyn-ChRmine-mScarlet-Kv2.1. All animal experiments were conducted in accordance with the approval of the Animal Care and Use Committee at the Shenzhen Institute of Advanced Technology, Chinese Academy of Sciences, and the Institutional Animal Care and Use Committee of PKU-Nanjing Institute of Translational Medicine (Approval ID: IACUC-2021-018).

### Neural and behavioral data collection

The collection of neural and behavioral data is based on the MouseVenue3D system. Four Intel RealSense D435 cameras are mounted orthogonally on four supporting pillars made of stainless steel. The distance between the nearest cameras is 85 cm. The cameras are adjusted to 45 cm off the ground to capture the whole view of the animal activities in the open field. Images were simultaneously recorded at 30 frames in 640×480 sizes by a PCI-E USB-3.0 data acquisition card and the PyRealsense2 Python camera interface package. The cameras are connected to a high-performance computer (i9 10900K, 128G RAM) equipped with a 512-gigabyte SSD and two 16-terabyte HDDs as an image acquisition platform. The computer also controls the camera calibration module.

The version of the miniature two-photon microscopy (mTPM) system is NUPERNOVA-600 (TRANSCEND VIVOSCOPE). The frame size is 512×512 and the frame rate is 4.84 Hz. The synchronization of 4 behavioral cameras and mTPM is based on the integrated synchronization module in mTPM. MouseVenue3D would send a TTL marker to mTPM every 30 frames. mTPM received the marker and output to the hardware time to a file. At the same time, mTPM writes the hardware time of every neural recording frame to another file. Later data alignment is based on these two timestamp files.

Two types of experiments were performed, including neural behavioral mapping and neural behavioral perturbation. In the neural behavioral mapping experiments, a subject mouse (implanted with an mTPM) and an object mouse (untreated) were allowed to interact freely in an open field for 15 minutes. During this period, we simultaneously recorded the subject’s mTPM neural activity and multi-view behavior videos of both animals. In the neural behavioral perturbation experiments, the subject and object mice first engaged in 15 minutes of free social interaction without optogenetic stimulation, during which mTPM neural activity and multi-view behavior were recorded. Subsequently, the optogenetic laser was switched on, and a second 15-minute session of simultaneous mTPM imaging and behavioral recording was performed. Because the fluorescence intensity of the perturbation experiment is weaker than that of mapping preparations, and ambient illumination can degrade imaging quality, we reduced the background lighting level during optogenetic experiments to ensure stable image quality. Within each experiment, the ambient illumination level was kept constant across pre-stimulation and stimulation sessions.

### The quality control of neural data

The neural activities are extracted by suite2p in the 4.84 Hz frame rate. Other parameters of suite2p are kept default. The extracted neural activities are upsampled using step interpolation to 30 frames per second to align with the behavior data. In order to eliminate the frequency artifact caused by the upsample, the upsampled neural activities are filtered by an equiripple lowpass filter with 2.0 passband frequency, 2.2 stopband frequency, and 20 density factor. Further, the filtered neural data *F* (*t*) are processed by the quality control step. We firstly extracted the running baseline *F*_0_ (*t*), which was estimated as *F*_0_ (*t*) = *F*_*s*_ (*t*) + *m*. *F*_*s*_ (*t*) is the smoothed fluorescence signal, calculated as the eighth percentile of *F* (*t*) within a ±15s moving window centered at time *t*. *m* is a small constant value calculated by *m* = min(*F*_*std*_ (*t*)) + 0.1*(min(*F*_*std*_ (*t*)) + max(*F*_*std*_ (*t*))). The neural activities finally expressed as the fractional change in the fluorescence with respect to baseline 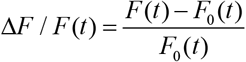. We manually checked that the baseline of Δ*F* /*F* is flatten but they existed noisy outliers. We set two hard thresholds *T*_min_ = −10 and *T*_max_ = 20 to exclude the neurons with high outliers.

Because optogenetic imaging exhibits a relatively low signal-to-noise ratio (SNR), we applied A fast blind zero-shot denoiser (Noise2Fast) to enhance image quality. Noise2Fast was used with its default parameters. To ensure a consistent noise baseline across conditions, every frame of both the mTPM imaging and the corresponding perturbation sessions in the same mouse was denoised using Noise2Fast. The denoised image sequences were then processed by Suite2p for motion correction, ROI segmentation, and fluorescence extraction.

To further improve the SNR of the extracted neural signals, we implemented a one-dimensional (1D) Unet following the strategy of DeepCAD-RT. The 1D Unet consisted of an encoder and decoder, each composed of four layers of 1D convolutional blocks. The model was trained in a self-supervised manner using the odd-indexed frames as input and the even-indexed frames as targets (and vice versa). Training was performed with a mini-batch size of 16, a maximum of 20 epochs, and a learning rate of 1×10^−4^. Because the predicted output sequence has half the temporal resolution of the original, we applied linear interpolation to restore the denoised signals to their original length. After that, the above upsampling, filtering, and baseline correction are applied to the denoised neural data.

### Pose estimation using Anti-Drift Pose Tracker (ADPT)

ADPT was trained on one NVIDIA GeForce RTX 3090 GPU. The definition of poses is the same as including “0: nose”, “1: left_ear”, “2: right_ear”, “3: neck”, “4: left_front_limb”, “5: right_front_limb”, “6: left_hind_limb”, “7: right_hind_limb”, “8: left_front_claw”, “9: right_front_claw”, “10: left_hind_claw”, “11: right_hind_claw”, “12: back”, “13: root_tail”, “14: mid_tail”, and “15: tip_tail”. The sleletons are definded including [0,1], [0,2], [1,2], [2,3], [1,3], [3,4], [3,5], [4,5], [4,6], [4,8], [5,7], [6,7], [5,9], [6,10], [7,11], [3,12], [4,12], [5,12], [6,12], [7,12], [13,12], [7,13], [6,13], [13,14], and [14,15]. Other parameters are set to the default.

We used the single animal pose estimation function of ADPT. We separately labeled 2615 frames of mice mounted FHIRM-TPM and 1852 frames of mice without neural recordings. Two ADPT models are trained separately, and the predictions of the poses of each mouse are concatenated together for later 3D reconstruction.

The 3D reconstruction of two mice in the same open field has two steps. The first step is the 2D to 3D projection. The basic projection formula between 2D points and 3D space points is as follows.

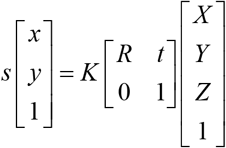

Here, *s* represents the scaling factor, *x* and *y* are the points in the image, *K* is the camera internal reference, *R* is the rotation matrix, *t* is the translation matrix, and *X*, *Y* and *Z* represent the coordinates of the 3D points. The camera calibration step gave *K*, *R*, and *t* of each camera. Combing with the 2D points [*x, y*,1]^*T*^ frame by frame, the 3D points [*X*,*Y*, *Z*,1]^*T*^ can be soloved.

The second step of 3D reconstruction is ground rotation. The 3D skeletons from the first step exist in the relative coordinates to the calibration checkboard. Because the calibration checkboard is not always parallel to the ground, the 3D skeletons need to be rotated to the ground coordinates. We used the 3D points of two mice’ hind paws to estimate the locations of the ground. We applied least square method to fit the plane of these 3D points as follows.

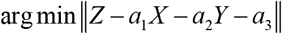

Here, *X*, *Y*, and *Z* are the coordinates of 3D points. *a*_1_ and *a*_2_ are the weights of *X* and *Y*. *a*_3_ is the intercede. So, the normal vector of the fitted plane is [*a*_1_, *a*_2,_, *a*_3_]. The ground plane is defined to be parallel to the ground so its normal vector is [0, 0, −1]. Then the cross vector is [−*a*_2_, *a*_1_, 0] and the rotation vector is 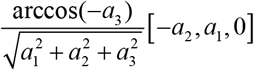. The rotated 3D skeletons [*X*_*r*_,*Y*_*r*_, *Z*_*r*_] can be calculated through Rodrigues formula *R*(·):

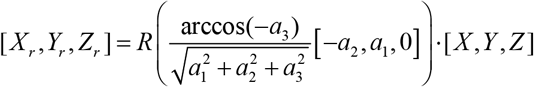

### Movement decomposition and identification using Behavior Atlas (BeA)

BeA was trained on a PC with an Intel i9-12900K CPU and 128 GB DDR4 RAM using the rotated 3D skeleton data. The size of the smooth window of BeA is 500 ms, and the smoother is a median filter. The body points of “12: back” and “13: root_tail” are used to align the direction of skeletons. The body sizes of each frame are corrected to 25-pixel-centered distributions according to the length between “12: back” and “13: root_tail”. “14: mid_tail” and “15: tip_tail” are not selected for later movement decomposition. The percentage of temporal dimensional reduction of the first layer is 5. The clustering number of the first layer is 3. The kernel of the second layer is Gaussian. The kernel size of the second layer is 30. The clustering number of the second layer is 24. The minimum and maximum time windows of the second layer are 500 and 2000 ms. The initialization method of the second layer is spectral clustering. Other BeA parameters are set to default. The movement decomposition is done one by one for each mouse in each 15-minute video.

Further, the decomposed movements are reclusted for phenotype identification. Pairwise dynamic time alignment kernel (DTAK) distances are calculated between all of the movement segmentations. The kernel size is 30. The DTAK distance matrix is embedded using uniform manifold approximation and projection (UMAP) with 199 n_neighbors and 0.05 min_dist. Hierarchical clustering with standard Euclidean distance and Ward linkage is used to cluster the UMAP embedding. The clustering number is 80. The recluster is separately done in the mice with or without FHIRM-TPM. In total, 160 clusters of movements are manually identified. The clusters with the same name are merged together. Finally, 49 movement categories are defined.

### Sequence extraction using Body Language-Bidirectional Encoder Representation from Transformers (BL-BERT)

BL-BERT was trained on one NVIDIA GeForce RTX 3090 GPU. 400,000 sequences are sampled from the movements for training. The length of each sequence is 120 movement modules. The percentage of masks is 0.2. The training batch size is 50. The number of epochs is 20. The initial learning rate is 1.0. BL-BERT is trained using a warm-up of 3000 iterations. The resampling time of the mask for prediction is 100. The 35 cluster number is selected according to the silhouette coefficient, based on the Hamming distance matrix across each pair of the sequences.

### The separation of social interaction episodes and post-interaction trials

According to our previous studies, the change point between close interaction and other behavior is about 5 cm of body distance, which can be converted to around 60 pixels using 0.88 cm/pixel^24^. Using 5 cm as the threshold between close interaction and other behavior, the full behavior data can be separated into dynamic segments. 5 s is chosen as the baseline of close interaction and the time after close interaction in the neural behavioral mapping data. The time length of the close interaction part is set to 2 s to 10 s to exclude the very short interaction and long-time stay close but no interaction. Using these criteria, 98 trials are separated from all 15 videos for further analysis. The trials are sorted according to their length for visualization.

In the perturbation experiments, we initially evaluated a 5 s baseline window. However, the neural principal component (PC) trajectories did not exhibit a clear or reproducible pattern under this baseline length. After systematic inspection, we found that shortening the baseline window enhanced the visibility of consistent neural patterns. This limitation is likely related to the character of PCA. The baseline dynamics differ substantially from those during social interaction episodes, and the relatively low signal-to-noise ratio of the perturbation recordings makes the extracted PCs more susceptible to baseline-induced variance. We therefore tested a shorter 2 s baseline, which consistently revealed a robust and reproducible PC pattern in the perturbation data. Based on this empirical optimization, a 2 s baseline window was used for all subsequent perturbation analyses. Correspondingly, the time length of the close interaction part is set to 2 s to 4 s to match the shorter baseline.

The post-interaction trials are separated using the same 5 cm distance threshold. Considering that the concerned data are the after-interaction, we chose 1 s close interaction as the baseline to keep as much data as possible. The time length of the after-interaction trials is from 10 s to 180 s, which firstly covers the baseline and after-length of close interaction trials, and secondly contains as many trials as possible. Using these criteria, 219 after-interaction trials are separated from all 15 videos for further analysis. The trials are sorted according to their length for visualization.

### Linear mapping using the PCs of neural activities to hierarchical behaviors

Before linear mapping, the neural activities are transformed using principal component analysis (PCA). The PCA is used in each of the trials above. The coefficients of PCA are normalized and aligned according to their length, one PC by one PC. Because the behavior has three hierarchies and their continuities are different, we applied two linear models to separately map neural PCs with continuous poses and discrete movements and sequences.

The generalized linear model (GLM) is used for linear mapping between neural PCs and poses because they are both continuous variables. The neural PCs of subject mice are separately mapped to the subject and object poses trial by trial using GLM in close interaction data. The discriminant analysis is used for linear mapping between neural PCs and movements and also between neural PCs and sequences because the output movement and sequence labels are discrete variables. The neural PCs of subject mice are separately mapped to the subject/object movements/sequences trial by trial. The mean mapping accuracy of all of them is 99.81±0.63%. The high mapping accuracy has verified that GLM and discriminant analysis can weight the combination of different neural PCs to the optimized fitting of poses, movements, and sequences.

### The parameter settings of the recurrent switching linear dynamic system (rSLDS)

We used rSLDS to describe the dynamics of the neural data in this paper. Before using rSLDS, the data are intercepted from the beginning of close interactions to keep each data having the same initial baseline. The threshold of distance to separate close interaction is 5 cm. The baseline of the beginning of close interactions is 15 s. Each behavior together with neural recording data is cut to 600 s from the start of the baseline to the end. Further, the data are separated into 3 groups for rSLDS modeling.

The first group is the close interaction group. This step is similar to the separation of close interaction trials. The minimum time of close interaction is 2 s, the baseline is 2 s, and the time after interaction is 2 s. The numbers of trials of S1, dmPFC, and M1 are separately 13, 17, and 25. The second group is the after-interaction group. This step is also similar to the separation of after-interaction trials. The minimum time of after-interaction is 5 s. The baseline is 1 s. The numbers of trials of S1, dmPFC, and M1 are separately 60, 80, and 79. The third group is the all-data group. Each trial is the 600 s data described before. After the separation of these 3 groups, their neural PCs are calculated trial by trial. The first 150 PCs of each trial are kept. The 150 neural PCs inside each brain area are concatenated for training rSLDS one by one brain area across 3 groups.

The rSLDS model consists of a discrete state *z*_*t*_ ∈{1,…, *K*}, a continuous latent state *x*_*t*_ ∈ ℝ^*D*^, and an observation 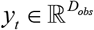. In this study, we set *K* = 2 (two latent dynamical regimes emerging from the concatenated PCA trajectories), *D* = 2 (for visualization and interpretability), and *D*_*obs*_ = 150 (the forst 150 PCs of neural activity). The generative process for each discrete state *k* is:

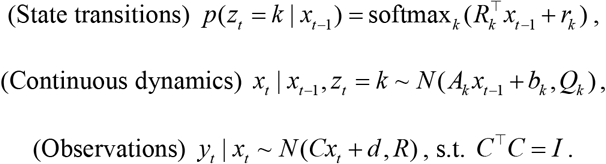

Recurrent transitions (transitions=“recurrent_only”) allow the switching probabilities to depend solely on the previous continuous latent state *x*_*t* −1_. State-specific dynamics use diagonal linear Gaussian matrices *A*_*k*_ (dynamics=“diagonal_gaussian”) to reduce overfitting and enable vector field interpretation. Orthogonal Gaussian emissions (emissions=“gaussian_orthog”, single_subspace=True) ensure that the neural observation space projects into a stable, interpretable low-dimensional subspace. Model parameters directly correspond to the script variables: *R* is transitions.Rs, *r* is transitions.r, *A*_*k*_ is dynamics.As, and *b*_*k*_ is dynamics.bs.

We fit a separate rSLDS model for each cortical region using Laplace-EM variational inference (method=“laplace_em”) with a structured mean-field posterior (variational_posterior=“structured_meanfield”). The initialization is random with a fixed seed for reproducibility. The iteration time is 1000. The optimization target is the evidence lower bound. The posterior outputs are the continuous latent means 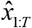 and the most likely discrete state sequence 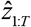 were extracted using 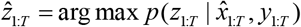.

To visualize the state-dependent flow fields, we constructed a 2D grid over the latent space and computed the one-step drift Δ*x* = (*A*_*k*_ *x* + *b*_*k*_) − *x*. Each grid point was assigned to its most likely discrete state via the recurrent transition function 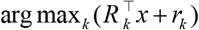. Vector fields show Δ*x* arrows within each state region. Plot limits were adapted to 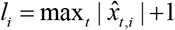 to show the full fields.

### Non-locomotion speed

Non-locomotion speed was used to quantify the animal’s movement when no net body displacement occurred (i.e., excluding walking or running). This metric captures subtle kinematic events such as micromovements, sniffing, turning-in-place, and other posture changes that do not alter the animal’s global position but reflect fine-scale motor dynamics. Let [*x*_*t*_, *y*_*t*_, *z*_*t*_] denote the 3D position of the animal at time *t*. The instantaneous speed is

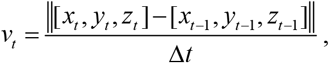

where Δ*t* is the interval time between two frames.

Non-locomotion speed is defined as

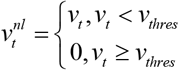

where *v*_*thres*_ is the threshold to separate non-locomotion speed and locomotion speed. We applied 150 pixels/s as *v*_*thres*_, empirically determined as approximately three times the mean baseline speed. Frames with instantaneous movement below this threshold were categorized as non-locomotion movements. The mean non-locomotion speed is then

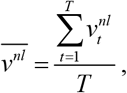

where *T* is the frame length of the total time.

### The mTPM illustration using max projection

To visualize the spatial activation of neurons in mTPM recordings, we generated maximum-intensity projection images from mTPM imaging stacks. For each imaging session, a mean reference frame was computed across the entire recording for each stack and subtracted from each frame to obtain a baseline-corrected signal. The resulting frames were normalized to the range [0,1] using linear rescaling. To suppress slow baseline drifts and emphasize transient activity, we applied a temporal rolling normalization using a sliding window of 100 frames. This rolling normalization enhanced local fluctuations while preserving spatial structure. Finally, for each stack, we computed a maximum-intensity projection across the temporal dimension:

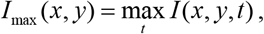

where *I* (*x, y,t*) denotes the normalized, rolling-corrected fluorescence signal at pixel (*x, y*) and time *t*. The resulting max-projection images provide a compact representation of the spatial trace of neural activity over the entire recording and were used for visualization of mTPM activity patterns.

### PC predictability

To quantify the dynamical predictability of each neural PC, we estimated its one-step temporal predictability using a minimal first-order autoregressive (AR(1)) model combined with K-fold cross-validation. For each PC time series 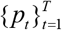, we constructed paired samples (*p*_*t*_, *p*_*t* +1_) and fit a linear autoregressive model of the form:

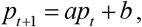

where *a* and *b* were estimated by ordinary least squares.

To obtain a robust and unbiased estimate of predictability, we performed k-fold cross-validation (k=30). In each fold, the model was trained on the training subset and evaluated on held-out samples. Prediction errors were accumulated across all folds. We quantified predictability using Root Mean Squared Error (RMSE) metrics:

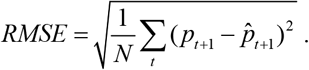

Lower RMSE indicates higher temporal predictability and stronger dynamical stability of the corresponding PC. This predictability metric captures the degree to which each neural PC follows a stable low-order dynamical rule, and was used to assess selective destabilization of high-dimensional components under optogenetic perturbation.

### Directional alignment analysis between neural predictability and behavior

To assess whether optogenetic-induced changes in neural dynamics were behaviorally relevant, we examined the directional consistency between changes in PC predictability and changes in non-locomotion speed across individual mice. For each mouse *i* and each PC *k*, we computed:

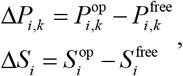

where *P* denotes PC predictability metric and *S* denotes the mean value of non-locomotion speed of each body point, under optogenetic (op) and control (free) conditions.

We focused on the directional relationship between neural destabilization and behavioral change caused by optogenetic perturbation. For each PC *k*, we counted across mice the number of cases in which the sign of predictability change was opposite to the sign of speed change:

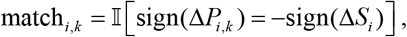

which reflects the hypothesis that increased neural unpredictability is associated with reduced non-locomotion speed, and vice versa. To avoid spurious contributions from near-zero fluctuations, mice with |Δ*P*_*i,k*_|< *ε*_*P*_ or |Δ*S*_*i*_|< *ε*_*S*_ were excluded (*ε* = 10^−6^).

For each PC, we performed a one-sided exact binomial test against the null hypothesis of random directional alignment (probability *p* = 0.5). The observed number of directionally consistent mice *m*_obs_ was compared to the binomial distribution:

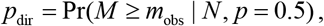

where *N* is the number of valid mice.

This test evaluates whether optogenetic-induced neural destabilization in a given PC is systematically aligned with no-locomotion speed changes across animals. PCs showing significant directional alignment indicate dimensions in which neural dynamical perturbations are kinematically relevant.

### Radius of rSLDS latent trajectories

To quantify the geometric dispersion of latent trajectories inferred by rSLDS, we measured the radial extent of each trajectory in the two-dimensional latent space. For each latent trajectory 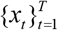, we first computed its centroid:

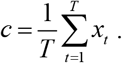

We then computed the instantaneous radius at each time point as the Euclidean distance from the centroid:

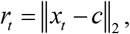

and reported the mean trajectory radius as:

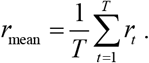

This mean radius summarizes how widely the latent trajectory disperses around its center; larger values indicate more spatially extended latent dynamics, whereas smaller values indicate more compact dynamics.

### Relative radial variability

Based on *r*_*t*_, the relative radial variability is calculated by

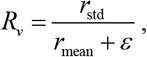

where *r*_std_ is the standard deviation. This dimensionless ratio measures the relative fluctuation of trajectory spread, independent of its absolute scale. Small *R*_*v*_ values indicate that the trajectory maintains a nearly constant radius, corresponding to geometrically stable and confined dynamics, whereas large *R*_*v*_ values indicate strong radial fluctuations, reflecting irregular, unstable, or intermittently expanding latent dynamics.

### Angular sweep score

To quantify the rotational structure of latent trajectories, we defined an angular sweep score that measures the degree of persistent angular progression around the trajectory centroid. For each 2D latent trajectory 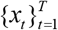, we first computed its centroid *c* as we introduced before. We then expressed each point in polar coordinates relative to the centroid and computed the instantaneous angular position:

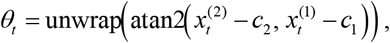

where unwrap(·) removes 2*π* discontinuities to obtain a continuous angular trajectory. We defined the angular increment as Δ*θ*_*t*_ = *θ*_*t*+1_ −*θ*_*t*_. The angular sweep score was then computed as:

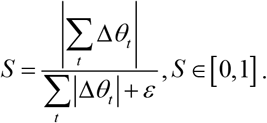

This score quantifies the net angular displacement normalized by the total angular path length. *S* ≈ 1 indicates persistent unidirectional rotation, corresponding to sustained cyclic or orbit-like dynamics. *S* ≈ 0 indicates frequent reversals of angular direction, corresponding to back-and-forth, non-rotational trajectories.

### Statistics

Before hypothesis testing, data were first tested for normality by the Shapiro-Wilk normality test and for homoscedasticity by the F test. For normally distributed data with homogeneous variances, parametric tests were used; otherwise, non-parametric tests were used. All of the ANOVA analyses are corrected by the recommended options of Prism 8.0. No data in this work are removed. All related data are included in the analysis.

## Data availability

We provide all model weights and the validation data used in this paper at

https://doi.org/10.6084/m9.figshare.30773648,

https://doi.org/10.6084/m9.figshare.30781931,

https://doi.org/10.6084/m9.figshare.30771338,

https://doi.org/10.6084/m9.figshare.30772454,

https://doi.org/10.6084/m9.figshare.30772685,

https://doi.org/10.6084/m9.figshare.30772766,

https://doi.org/10.6084/m9.figshare.30773072.

They also contain the necessary intermediate data for the validation and visualization of this study. This paper can be replicated using the downloadable data and supplied code.

## Code availability

All codes to reproduce this work is at https://github.com/YNCris/lmt_code.

## Acknowledgements

This work was supported in part by STI2030-Major Projects grant 2021ZD0202205 (HC), STI2030-Major Projects grant 2021ZD0203900 (PW), National Natural Science Foundation of China grant T2394533 (PW), Jiangsu Provincial Major Science and Technology Special Project grant BG2025040 (PW), Jiangsu Provincial Advanced Technology Research and Development Program grant BF2025060 (PW), and Start-up Research Fund of Southeast University grant RF1028625096 (PW).

## Author contributions

Conceptualization: YH, QL, SL, HC, PW

Methodology: YH, PW

Investigation: YH

Visualization: YH, QL, SL, HC, PW

Funding acquisition: HC, PW

Project administration: QL, SL, PW

Supervision: YH, QL, SL, HC, PW

Writing – original draft: YH

Writing – review & editing: YH, QL, SL, PW

